# Carbon inputs from riparian vegetation limit oxidation of physically-bound organic carbon via biochemical and thermodynamic processes

**DOI:** 10.1101/105486

**Authors:** Emily B. Graham, Malak M. Tfaily, Alex R. Crump, Amy E. Goldman, Lisa Bramer, Evan Arntzen, Elvira Romero, C. Tom Resch, David W. Kennedy, James C. Stegen

## Abstract

In light of increasing terrestrial carbon (C) transport across aquatic boundaries, the mechanisms governing organic carbon (OC) oxidation along terrestrial-aquatic interfaces are crucial to future climate predictions. Here, we investigate the biochemistry, metabolic pathways, and thermodynamics corresponding to OC oxidation in the Columbia River corridor using ultra-high resolution C characterization. We leverage natural vegetative differences to encompass variation in terrestrial C inputs. Our results suggest that decreases in terrestrial C deposition associated with diminished riparian vegetation induce oxidation of physically -bound OC. We also find that contrasting metabolic pathways oxidize OC in the presence and absence of vegetation and—in direct conflict with the ‘priming’ concept—that inputs of water-soluble and thermodynamically favorable terrestrial OC protects bound-OC from oxidation. In both environments, the most thermodynamically favorable compounds appear to be preferentially oxidized regardless of which OC pool microbiomes metabolize. In turn, we suggest that the extent of riparian vegetation causes sediment microbiomes to locally adapt to oxidize a particular pool of OC, but that common thermodynamic principles govern the oxidation of each pool (*i.e*., water-soluble or physically-bound). Finally, we propose a mechanistic conceptualization of OC oxidation along terrestrial-aquatic interfaces that can be used to model heterogeneous patterns of OC loss under changing land cover distributions.

**Key points:**

- Riparian vegetation protects bound-OC stocks
- Biochemical processes associated with OC oxidation vary with vegetation conditions
- Common thermodynamic principles underlie OC oxidation regardless of vegetation conditions

## 1. Introduction

Soils and nearshore sediments comprise a carbon (C) reservoir that is 3.2 times larger than the atmospheric C pool [*Burd et al.*, 2016], yet Earth System Models (ESMs) struggle to integrate mechanisms of OC oxidation in these environments into predictions of atmospheric carbon dioxide concentrations [*Todd-Brown et al.*, 2013; *Wieder et al.*, 2013; *Wieder et al.*, 2015]. In particular, OC oxidation in nearshore habitats constitutes a significant uncertainty in atmospheric C flux [*Aalto et al.*, 2003; *Battin et al.*, 2009] and knowledge about C cycling along these transitional ecosystems is necessary to accurately predict global C cycling [*Burd et al.*, 2016]. Terrestrial C inputs into aquatic systems have nearly doubled since pre-industrial times; an estimated 2.9 Pg C now crosses terrestrial-aquatic interfaces annually (vs. 0.9 Pg C yr^-1^ stored within forested ecosystems) [*Battin et al.*, 2008; *Regnier et al.*, 2013]. The magnitude of this flux has garnered significant recent attention [*Battin et al.*, 2008; *Battin et al.*, 2009; *Regnier et al.*, 2013], yet the biochemical, metabolic, and thermodynamic mechanisms governing OC oxidation along terrestrial-aquatic interfaces remain a crucial uncertainty in climate predictions. New molecular techniques are providing insight into OC dynamics [*Mason et al*, 2016; *Malak M Tfaily et al.*, 2015; *M.M. Tfaily et al.*, 2017], but we still lack an understanding of why some OC remains stabilized for millennia whereas other OC is rapidly oxidized [*Schmidt et al.*, 2011].

The ability of microorganisms to oxidize complex OC is an important constraint on C cycling because OC is a mixture of compounds with different propensities for biotic oxidation [*Hedges and Oades*, 1997; *Hedges et al.*, 2000]. Within terrestrial research, OC oxidation is often framed within the concept of ‘priming,’ whereby microbial oxidation of chemically-complex, less bioavailable OC is fueled by the addition of more bioavailable and thermodynamically favorable OC compounds [*Kuzyakov*, 2010]. However, the applicability of priming in aquatic environments is unclear [*Bengtsson et al.*, 2014; *Bianchi*, 2011; *Guenet et al.*, 2010]. Aquatic systems, and in particular nearshore environments, frequently experience mixing of terrestrial and aquatic C sources with distinct chemical character, providing a theoretical basis for priming expectations [*Bengtsson et al.*, 2014; *Guenet et al.*, 2010]. Consistent with priming, *Guenet et al.* [2010] proposed that this mixing generates “hotspots” or “hot moments” of biological activity facilitated by complementary C resources. Alternatively, OC stabilization in sediments is tightly linked to organomineral interactions, which provide physical protection from extracellular enzyme activity [*Hedges and Keil*, 1995; *Hunter et al.*, 2016; *Rothman and Forney*, 2007], and the strength of these interactions may override any influence of priming. Early investigations of priming effects in aquatic systems have been inconclusive, with evidence both for [*Dorado-García et al.*, 2016] and against [*Bengtsson et al.*, 2014; *Catalán et al.*, 2015] priming mechanisms.

Several new perspectives have attempted to move beyond frameworks, such as priming, that depend on strict chemical definitions to predict OC oxidation [*Burd et al.*, 2016; *Cotrufo et al.*, 2013; *Lehmann and Kleber*, 2015]. Recent work proposes that the probability of OC oxidation is related to a spectrum of chemical properties and that even very complex OC can be oxidized when more thermodynamically favorable OC is depleted or isolated from microorganisms. For example, *Lehmann and Kleber* [2015] have proposed a ‘soil continuum hypothesis’ whereby OC is a gradient of continuingly decomposing compounds that are variably accessible for biotic oxidation, with no notion of chemically labile versus recalcitrant compounds. Similarly, *Burd et al.* [2016] have suggested that OC oxidation is a ‘logistical problem’ involving the ability of microorganisms to access and metabolize compounds. Both concepts capture the emerging belief that chemically-complex, less thermodynamically favorable OC can be oxidized when more favorable compounds are inaccessible.

Here, we address a critical knowledge gap in predicting the global C balance [*Aalto et al.*, 2003; *Battin et al.*, 2009; *Burd et al.*, 2016; *Regnier et al.*, 2013]—mechanisms governing OC oxidation along terrestrial-aquatic interfaces. Specifically, we investigate the biochemistry, microbial metabolism, and thermodynamics of OC oxidation in nearshore water-soluble and physically-bound (*i.e*., mineral and microbial) OC pools along a freshwater terrestrial-aquatic interface. We leverage natural variation in riparian vegetation along the Columbia River in Eastern Washington State, the largest river in the U.S. west of the Continental Divide [*Ebel et al.*, 1989; *Moser et al.*, 2003], to examine these mechanisms in the context of spatial variation in terrestrial C deposition. Consistent with the priming paradigm, we hypothesize that (a) C deposition associated with riparian vegetation increases total aerobic metabolism and enhances oxidation of bound-OC stocks, while (b) areas without riparian vegetation foster lower rates of aerobic metabolism with minimal oxidation of bound-OC.

## 2. Materials and Methods

### 2.1. Site Description

This study was conducted along the Columbia River shoreline within the Hanford Site 300 Area (approximately 46° 22’ 15.80”N, 119° 16’ 31.52”W) in eastern Washington State [*Graham et al.*, 2016a; 2017; *Slater et al.*, 2010; *Zachara et al.*, 2013]. The Columbia River experiences shoreline geographic variation in vegetation patterns, substrate geochemistry, and microbiome composition [*Arntzen et al.*, 2006; *Lin et al.*, 2012; *Peterson and Connelly*, 2004; *Slater et al.*, 2010; *James C Stegen et al.*, 2016; *James C. Stegen et al.*, 2012; *Zachara et al.*, 2013]. Accordingly, the Hanford Reach of the Columbia River embodies an ideal natural system in which to examine heterogeneity of terrestrial OC inputs and subsequent OC oxidation mechanisms.

Liquid N_2_-frozen sediment profiles (0-60 cm) were collected along two shoreline transects with or without riparian vegetation (hereafter, V and NV for ‘vegetated’ and ‘not vegetated’, Table 1, Figure 1) perpendicular to the Columbia River in March 2015, separated by a distance of ~170m. V was characterized by a moderately sloping scour zone, small boulders, and a closed canopy of woody perennials *Morus rubra* (Red Mulberry) and *Ulmus rubra* (Slippery Elm). The highest elevation (upper bank) samples were collected within the rooting zone. In contrast, NV was characterized by a gradually sloping scour zone, cobbled armor layer, and minimal vegetation. We collected profiles at three locations in each transect with 5m spacing within a spatial domain of ~175 × 10m. In each transect, the lower bank profile was located at ~0.5m (vertical distance) below the midpoint, which was in turn ~0.5m below the upper bank profile (approximately 10m horizontal distance from upper to lower profiles). All cores were collected (see below) during conditions in which the sediments were fully saturated, either being underwater at the time of coring or recently underwater prior to coring. Each profile was sectioned into 10-cm intervals from 0-60cm. Because OC composition (see below for Methods) did not differ across upper (0-10cm) to lower (50-60cm) sections in each profile, each 10-cm section was used as a replicate sample to provide sufficient sample size (n >15 at each transect) for comparisons between the vegetated and non-vegetated transect.

**Table 1.** Acronyms and abbreviations used in this paper.

| Abbreviation/Acronym | Description |
| --- | --- |
| V | Transect with dense riparian vegetation (i.e., 'vegetated') |
| NV | Transect with sparse riparian vegetation (i.e., 'not vegetated') |
| V-W | Transect V, water extraction (water-soluble OC) |
| V-B | Transect V, chloroform extraction (bound-OC) |
| NV-W | Transect NV, water extraction (water-soluble OC) |
| NV-B | Transect NV, chloroform extraction (bound-OC) |
| C | Carbon |
| OC | Organic carbon |
| FTICR-MS | Fourier transform ion cyclotron resonance mass spectrometry |
| KEGG | Kyoto Encyclopedia of Genes and Genomes |
| $\Delta G^0_{\text{Cox}}$ | Gibbs free energy of C oxidation |

**Figure 1.**
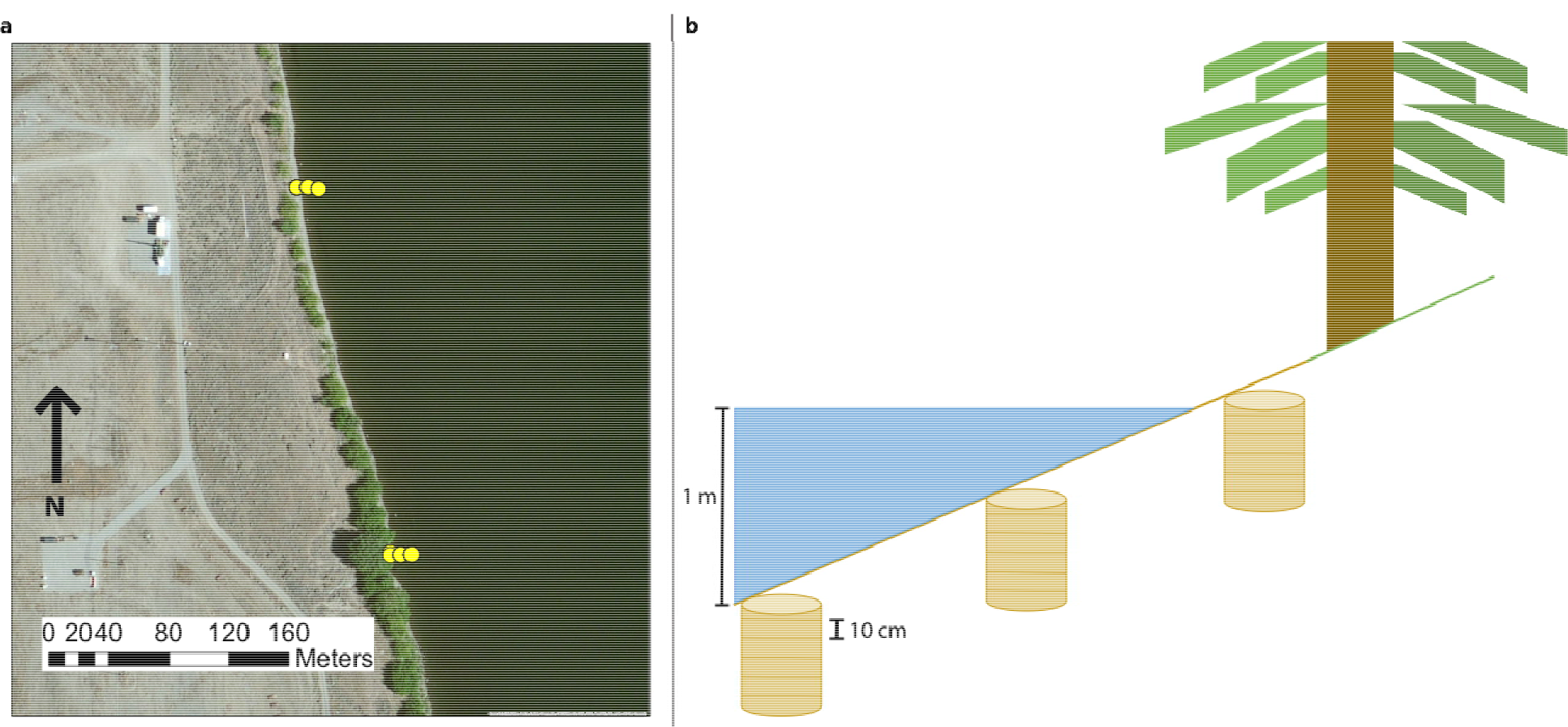
Schematic of Sampling Design. Samples were taken from three cores across an elevation gradient in a transect with or without dense vegetation. Panel (a) displays a satellite image of core locations. Cores without dense vegetation were obtained from the northern transect. Panel (b) shows a schematic of our sampling design. Within each core, samples were taken in 10-cm intervals from 0 to 60 cm. We extracted OC pools from each sample using a sequential extraction protocol (H_2_O, MeOH, and CHCl_3_) to target water-soluble and bound-OC pools.

### 2.2. Sample Collection

Liquid N_2_-frozen sediment profiles were collected as outlined in *Moser et al.* [2003] using a method developed by *Lotspeich and Reed* [1980] and modified by *Rood and Church* [1994]. A pointed stainless steel tube (152 cm length, 3.3 cm outside diameter, 2.4 cm inside diameter) was driven into the river bed to a depth of ~60cm. Liquid N_2_ was poured down the tube for ~15 minutes, until a sufficient quantity of material had frozen to the outside of the rod. The rod and attached material were removed from the riverbed with a chain hoist suspended beneath a tripod. Profiles were placed over an aluminum foil lined cooler containing dry ice. Frozen material was removed with a mallet. The material was then wrapped in the foil and transported on dry ice to storage at -80°C. In the lab, profiles were sectioned into 10cm depth intervals from 0-60cm in an anaerobic glove bag (n = 6 per profile, except for the lowest elevation in the non-vegetated transect which was sectioned only from 30-60cm due to low recovery yield from 0-30cm; total n = 33).

### 2.3. Physicochemistry

Details concerning physicochemical assays are provided in the Supporting Information. Briefly, we determined the particle distribution of sediments by separating size fractions via sieving; total nitrogen, sulfur, and carbon content were determined using an Elementar vario EL cube (Elementar Co.Germany); 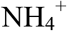 was extracted with KCl and measured with Hach Kit 2604545 (Hach, Loveland, Co); Fe(II) content was measured with a ferrozine assay; and all other ion concentrations were measured by inductively coupled plasma mass spectrometry (ICP-MS) on HCl extractions. Aerobic metabolism was determined with a resazurin reduction assay, modified from *Haggerty et al.* [2009].

### 2.4. Fourier transform ion cyclotron resonance mass spectrometry solvent extraction and data acquisition

We leveraged state of science chemical extraction protocols combined with Electrospray ionization (ESI) and Fourier transform ion cyclotron resonance (FTICR) mass spectrometry (MS) to infer differences in OC character among our samples. Previously, *Tfaily et al.* [2015; 2017] demonstrated the optimization of OC characterization from soils and sediments by sequential extraction with polar and non-polar solvents tailored to the sample set of interest. Tfaily et al.’s extraction procedures have been coupled to ESI FTICR-MS to distinguish OC pools among ecosystems and soil types [*Tfaily et al.*, 2015; *Tfaily et al.*, 2017] as well as to provide information on the metabolism of distinct OC pools among samples within a single environment [*Bailey et al.*, 2017]. Other common OC characterization methods such as nuclear magnetic resonance spectroscopy (NMR), Fourier transform infrared spectroscopy (FTIR), and gas chromatography MS only analyze a limited number of compound classes [*Kögel-Knabner*, 2002; *Kögel-Knabner*, 2000]. In contrast, ESI FTICR-MS introduces intact organic molecules into the MS without fragmentation and allows for the detection of a wide range of chemical compounds [*Tfaily et al.*, 2015; *Tfaily et al.*, 2017]. The use of 12 Tesla (T) FTICR-MS offers high mass resolving power (>1M) and mass measurement accuracy (<1 ppm), and while nascent in its application within complex environmental systems, it has emerged as a robust method for determining OC chemistry of natural organic matter [*Kim et al.*, 2003; *Koch et al.*, 2005; *Tfaily et al.*, 2011; *Tremblay et al.*, 2007]. Moreover, *Tfaily et al.* [2015; 2017] demonstrated that sequential extraction with targeted solvents can preferentially select OC pools with differing chemical character (*e.g*., lipid-like vs. carbohydrate-like).

Here, we used three solvents with different polarities—water (H_2_O), methanol (CH_3_OH, hereafter “MeOH”) and chloroform (CHCl_3_)—to sequentially extract a large diversity of organic compounds from samples, according to *Tfaily et al.* [2015; 2017]. Water extractions were performed first, followed by MeOH and then CHCl_3_. Previous work has shown that each solvent is selective towards specific types of compounds [*Tfaily et al.*, 2015]. Water is a polar solvent with a selection bias for carbohydrates with high O/C ratios, amino-sugars, and other labile polar compounds [*Tfaily et al.*, 2015]; and, as nearshore environments frequently experience wetting, water extractions represent an estimation of readily accessible OC compounds in these environments. Conversely, CHCl3 is selective for non-polar lipids associated with mineral interactions and cellular membranes (*i.e*., physically-bound OC) [*Tfaily et al.*, 2015]. Because MeOH has a polarity in between that of water and CHCl_3_, it extracts both water-soluble and bound-OC pools (*i.e*., a mix of compounds that water and CHCl_3_ extract), and *Tfaily et al.* [2015] demonstrated compositional overlap between water-soluble and MeOH extracted OC pools. In this study, we are interested in the differences in OC composition between pure water-soluble and bound-OC pools, and we focus our discussion on H_2_O- and CHCl_3_-extractions only. We use H_2_O- and CHCl_3_-extracted OC as proxies for readily bioavailable (*i.e*., weakly bound) vs. less bioavailable (*i.e*., mineral- and microbial-bound) pools, respectively.

Extracts were prepared by adding 1 ml of solvent to 100 mg bulk sediment and shaking in 2 mL capped glass vials for two hours on an Eppendorf Thermomixer. Samples were removed from the shaker and left to stand before spinning down and pulling off the supernatant to stop the extraction. The residual sediment was dried with nitrogen gas to remove any remaining solvent, and then the next solvent was added. The CHCl_3_ and H_2_O extracts were diluted in MeOH to improve ESI efficiency. *Tfaily et al.* [2015] estimated the OC extraction efficiency to be ~15%. *Tfaily et al.* [2015] have previously demonstrated extraction efficiencies as low as 2% to be representative of OC pool composition. We further note that numerous studies have established FTICR-MS as a robust method for distinguishing compositional differences among OC pools [*Herzsprung et al.*, 2017; *Kellerman et al.*, 2015; *Rossel et al.*, 2016; *Ward and Cory*, 2015; *Zhang et al.*, 2016].

Ultra-high resolution mass spectrometry of the three different extracts from each sample was carried out using a 12 Tesla Bruker SolariX FTICR-MS located at the Environmental Molecular Sciences Laboratory (EMSL) in Richland, WA, USA. As per *Tfaily at al.* [2017], we performed weekly calibration using a tuning solution containing C_2_F_3_O2, C_6_HF_9_N_3_O, C_12_HF_21_N_3_O, C_20_H_18_F_27_N_3_O_8_P_3_, and C_26_H_18_F_39_N_3_O_8_P_3_ with m/z ranging from 112 to 1333 (Agilent Technologies, Santa Clara, CA USA). Instrument settings were optimized using Suwannee River Fulvic Acid (IHSS). The instrument was flushed between samples using a mixture of water and methanol. Blanks were analyzed at the beginning and the end of the day to monitor for background contaminants.

The extracts were injected directly into the mass spectrometer, and the ion accumulation time was optimized for all samples to account for differences in OC concentration. The ion accumulation time ranged between 0.5 and 1s. A standard Bruker electrospray ionization (ESI) source was used to generate negatively charged molecular ions. Samples were introduced to the ESI source equipped with a fused silica tube (30 μm i.d.) through an Agilent 1200 series pump (Agilent Technologies) at a flow rate of 3.0 μL min^-1^. Experimental conditions were as follows: needle voltage, +4.4 kV; Q1 set to 50 *m*/*z*, and the heated resistively coated glass capillary operated at 180 °C. One hundred forty-four individual scans were averaged for each sample and internally calibrated using an organic matter homologous series separated by 14 Da (-CH_2_ groups). The mass measurement accuracy was less than 1 ppm for singly charged ions across a broad *m*/*z* range (100-1200 *m*/*z*). The mass resolution was ~ 350K at 339 m/z.

### 2.5 FTICR-MS data processing

A depiction of our FTICR-MS data processing pipeline is presented in Figure 2 and described in the following sections.

**Figure 2.**
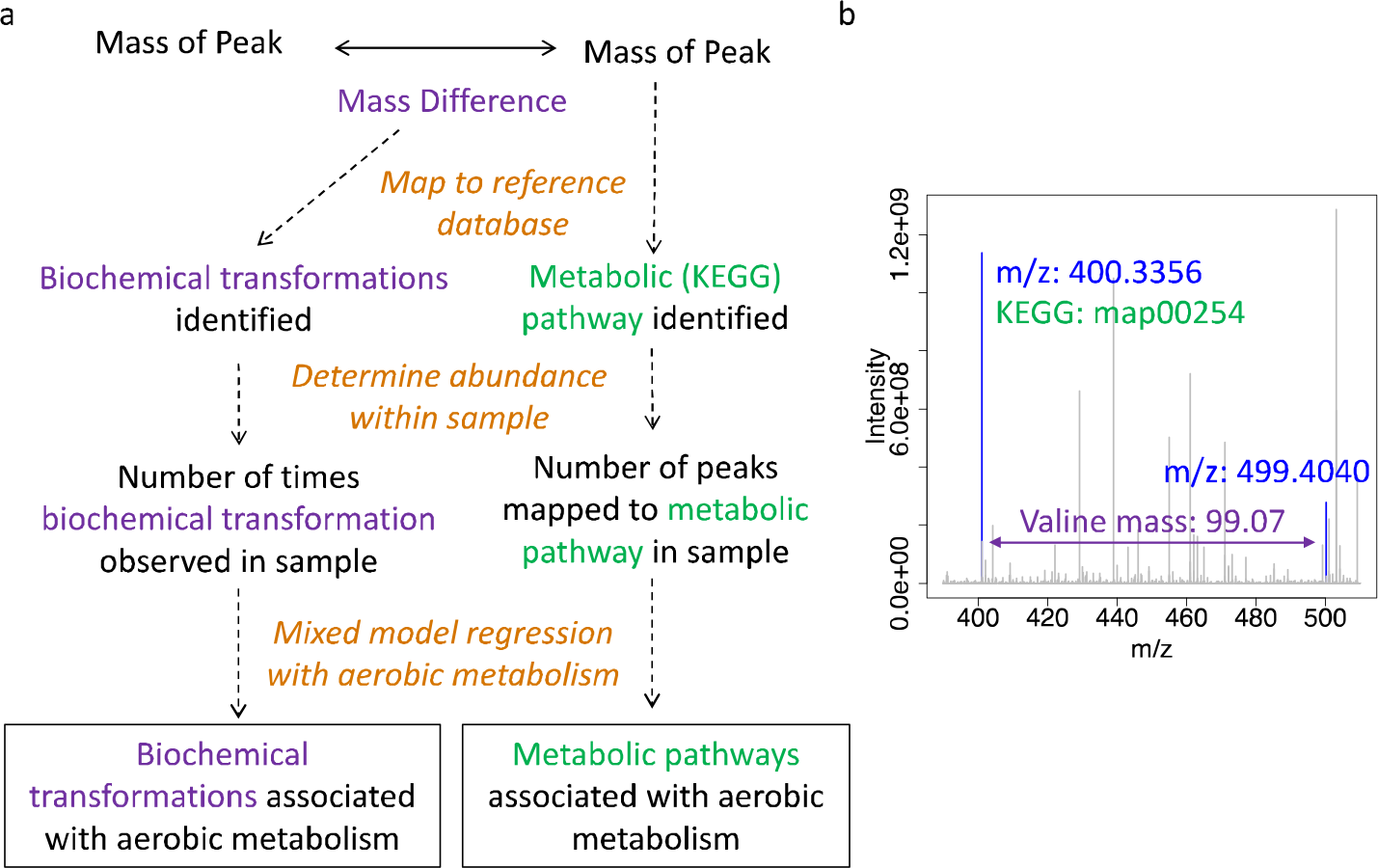
Methodology for inferring biochemical transformations and metabolic pathways. Panel (a) depicts our workflow for analyzing biochemical transformations and metabolic pathways. Biochemical OC transformations (purple) were identified by mapping mass differences in pairwise m/z peak comparisons to a set of 92 known masses transferred in common biochemical transformations (*e.g*., glucose, amines). Metabolic pathways (green) were identified by mapping all chemical formula assigned to m/z peaks to the KEGG database. Within each sample, the abundance of each biochemical transformation and the number of peaks mapping to each metabolic pathway were then correlated to aerobic metabolism to garner insights into OC oxidation processes. Panel (b) displays an example portion of our FTICR-MS spectra overlain with peak assignments (blue), a biochemical transformation (mass difference between peaks, denoted in purple), and a metabolic pathway (associated with the left-hand peak, denoted in green).

#### 2.5.1 Chemical formulae assignment

BrukerDaltonik software version 4.2 was used to convert raw spectra to a list of m/z values applying FTMS peak picker module with a signal-to-noise ratio (S/N) threshold set to 7 and absolute intensity threshold to the default value of 100. Putative chemical formulae were then assigned using in-house software following the Compound Identification Algorithm (CIA), proposed by *Kujawinski and Behn* [2006], modified by *Minor et al.* [2012], and previously described in *Tfaily et al.* [2017]. Chemical formulae were assigned based on the following criteria: S/N >7, and mass measurement error <1 ppm, taking into consideration the presence of C, H, O, N, S and P and excluding other elements. To ensure consistent formula assignment, we aligned all sample peak lists for the entire dataset to each other in order to facilitate consistent peak assignments and eliminate possible mass shifts that would impact formula assignment. We implemented the following rules to further ensure consistent formula assignment: (1) consistently pick the formula with the lowest error and with the lowest number of heteroatoms and (2) assignment of one phosphorus atom requires the presence of at least four oxygen atoms.

#### 2.5.2 Identification of putative biochemical transformations using FTICR-MS

To identify potential biochemical transformations, we followed the procedure detailed by *Breitling et al.* [2006] and employed by *Bailey et al.* [2017]. In essence, the mass difference between m/z peaks extracted from each spectrum with S/N>7 were compared to commonly observed mass differences associated with biochemical transformations. All possible pairwise mass differences were calculated within each extraction type for each sample, and differences (within 1ppm) were matched to a list of 92 common biochemical transformations (*e.g*., gain or loss of amino groups or sugars, Table S1). For example, a mass difference of 99.07 corresponds to a gain or loss of the amino acid valine, while a difference of 179.06 corresponds to the gain or loss of a glucose molecule. Pairs of peaks with a mass difference within 1 ppm of our transformation list were considered to be related by the corresponding compound. This approach is feasible with FTICR-MS data because the set of peaks in each sample are related by measureable and clearly defined mass differences corresponding to gains and losses of compounds. It has been previously used by *Bailey et al.* [2017] to demonstrate differences in biochemical transformations among soils incubated with different microbial inocula and among pore size classes in complex soil matrices.

#### 2.5.3 Identification of putative microbial metabolic pathways using FTICR-MS

Additionally, a set of putative microbial metabolic pathways in each sample can be identified by locating chemical formulae assigned to m/z’s within metabolic pathways defined in the Kyoto Encyclopedia of Genes and Genomes (KEGG, Release, 80.0, http://www.kegg.jp) [*Kanehisa and Goto*, 2000]. Chemical formulae were mapped to KEGG pathways using an in-house software to detect all KEGG pathways containing a giving formula. For example, a peak with a mass of 400.3356 was assigned formula C_20_H_16_O_9_ and mapped to KEGG pathway ‘map00254’ (Aflatoxin biosynthesis) which contains C_20_H_16_O_9_ as an intermediate. While only a subset of compounds detected by FTICR-MS are defined within the KEGG database (*i.e*., peaks must be assigned a chemical formula and that chemical formula must be present in a KEGG pathway), we found 415 unique peaks that were assigned putative molecular formulae *and* that corresponded to compounds present in KEGG pathways. Additionally, we defined assignments at the pathway level (*i.e*., by “map” number) instead of using enzyme level classification (*i.e*., EC number) in order to aggregate compounds found within the same pathways. This was done to facilitate functional interpretation.

Although we acknowledge our results do not represent a comprehensive analysis of all microbial metabolic pathways present in a sample, we assume that KEGG pathways containing more peaks detected by FTICR-MS within a sample are more likely to be active than those with fewer mapped peaks. We further reduced possible random matches by assessing correlations with aerobic metabolisms as described in the ‘Statistical Methods’ section below, and we compare results across samples to yield insight into microbial pathways in each sample beyond that which can be garnered from biochemical transformations. The results are, however, conceptually congruent with those derived from the biochemical transformation analyses described in the preceding sub-section. The KEGG pathway and transformation analyses are independent of each other, yet provided consistent insights, and thus together they provide greater confidence in our interpretations.

### 2.6 Statistical Methods

All statistical analyses were conducted using R software (https://www.r-project.org/). FTICR m/z intensities were converted into presence/absence data prior to analysis because differences in m/z intensity are influenced by ionization efficiency as well as relative abundance [*Kujawinski and Behn*, 2006; *Minor et al.*, 2012; *Tfaily et al.*, 2015; *Tfaily et al.*, 2017].

To examine differences in OC composition between transects, we used the ‘vegan’ package to construct a Sorenson dissimilarity matrix for all m/z’s identified (*i.e*., we included peaks with or without assigned formula) within each OC pool (water-soluble or physicallybound). Differences between vegetation states (*i.e*., V vs. NV) were tested with PERMANOVA with permutations stratified by depth to account for non-independence among samples from the same core (999 permutations, ‘vegan’) and visualized using Non-metric Multidimensional Scaling (NMDS, ‘vegan’). One sample (NV, upper profile, depth 30-40cm) was removed due to peak interference during FTICR-MS, and three samples (NV, middle profile, depths 00-10cm, 10-20cm, 20-30cm) were excluded, because we were unable to collect sufficient sample mass for all analyses.

To reveal transformations associated with aerobic metabolism and to study differences in those transformations across vegetation states, we determined the number of times a given transformation occurred within each OC pool in each sample. Specifically, for each of the 92 compounds in our set of biochemical transformations, we counted the number of times in each sample that transformation was observed to yield an estimate of the prevalence or ‘abundance’ of each transformation in each sample. We correlated these abundance estimates to rates of metabolism using linear mixed models in which transformations were a fixed effect and depth was a random effect. Mixed models were compared to null expectations (*i.e*., models including only random effects) with ANOVA to determine significance. Significant relationships were interpreted as biochemical transformations possibly associated with biotic OC oxidation. To evaluate how transformations associated with OC oxidation varied across vegetation states, we used the abundances of those transformations across all samples to calculate Bray-Curtis dissimilarity. Resulting Bray-Curtis dissimilarities were used to visualize multivariate differences among samples using non-metric Multidimensional Scaling (NMDS, ‘vegan’), and we statistically evaluated separation between vegetation states with PERMANOVA (999 permutations stratified by depth, ‘vegan’). We refer to H_2_O- and CHCl_3_-soluble OC pools at V and NV, respectively, as V-W (‘vegetated water’), V-B (‘vegetated bound’), NV-W (‘not vegetated water’), and NV-B (‘not vegetated bound’) for the remainder of the manuscript.

Similar to our analyses of biochemical transformations, we found the number of m/z’s that mapped to a given KEGG pathway. We make the assumption that pathways with more m/z’s mapped to them have a higher probability of actively contributing to biogeochemical function. To identify which pathways were most likely to contribute to aerobic metabolism, we tested the relationship between the number of m/z’s mapped to a given KEGG pathway within each sample and aerobic metabolism using linear mixed models with depth as a random variable as per our analyses of biochemical transformations. Pathways with significant relationships were interpreted as influencing OC oxidation, and the following analysis was conducted only with those pathways. To estimate relative abundances, the number of peaks mapping to each KEGG pathway in a sample was normalized by the total number of peaks mapping to any KEGG pathway (within the sample) that was significantly related to aerobic metabolism. To reveal groups of pathways co-varying with each other across vegetation states and OC pools, we statistically clustered pathways that correlated with aerobic metabolism. Clustering was based on pathway relative abundances in each vegetation state and pool type. Clusters were determined using the ‘hclust’ algorithm in R with the ‘complete linkage’ clustering method and visualized using the ‘pheatmap’ package.

Finally, we examined associations between aerobic metabolism and OC thermodynamics by calculating the Gibbs Free Energy of OC oxidation under standard conditions (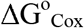) from the Nominal Oxidation State of Carbon (NOSC) as per *LaRowe and Van Cappellen* [2011]. NOSC was calculated from the number of electrons transferred in OC oxidation half reactions and is defined by the equation:

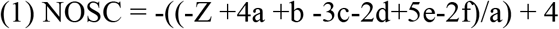

where a, b, c, d, e, and f are, respectively, the numbers of C, H, N, O, P, S atoms in a given organic molecule and Z is net charge of the organic molecule (assumed to be 1). In turn, 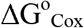 was estimated from NOSC following *LaRowe and Van Cappellen* [2011]:

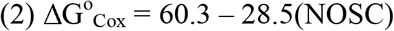

Values of 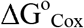 are generally positive, indicating that OC oxidation must be coupled to the reduction of a terminal electron acceptor. While 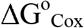 varies according to the availability of terminal electron acceptors, our system is primarily oxic, allowing us to infer oxygen as the primary electron acceptor in most reactions and make direct comparisons across samples. Additionally, though the exact calculation of 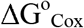 necessitates an accurate quantification of all species involved in every chemical reaction in a sample, the use of NOSC as a practical basis for determining 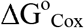 has been validated [*Arndt et al.*, 2013; *LaRowe and Van Cappellen*, 2011].

Here, we assessed relationships between aerobic metabolism and 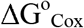 of OC compounds identified in each OC pool (determined by FTICR-MS analysis) using linear mixed models in each vegetation state, in which aerobic metabolism was a fixed effect, depth was a random effect, and average 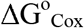 of all m/z’s with assigned formula was the dependent variable.

We note that we could not account for possible non-independence between the 3 cores obtained within each transect. Given one replicate per depth per core within each transect, core identity cannot be accounted for with mixed models, due to identifiability issues in model estimation, once accounting for depth. However, we emphasize that the locations within each transect were selected to represent a broad range of conditions within each vegetation state. That is, higher elevation cores were located near vegetation (vegetated state) or in a mostly barren cobbled bank (unvegetated state) in locations that submerge seasonally; lower elevation cores were in locations that are usually submerged below river water and the middle elevation represented a transitional location (Figure 1). The three positions within each transect therefore sampled a broad range of conditions with the potential to cause variation in OC composition and cycling. This sampling design enabled a regression-based analysis.

We originally hypothesized that OC composition/cycling would vary continuously across the broad range of sampled conditions. Our analyses, however, showed that for some key aspects of the data, the two sites grouped together instead of showing continuous variation. That is, despite within-transect variation in the physical environment, we observed coherence in OC composition/cycling within each transect. In turn, we provisionally infer that the presence or absence of riparian vegetation strongly constrained OC composition/cycling within the hyporheic zone of our system. The inference is made possible by the two transects being near each other such that most other features of the study system were similar between the two transects (e.g., aqueous chemistries, hydrology, mineralogy, temperature, light cycles, etc.). The major feature distinguishing the two transects was the presence or absence of riparian vegetation. However, as noted above, we consider our inferences to be foundational for future studies because we did not sampling multiple plots within each vegetated state.

## 3. Results and Discussion

### 3.1. Shifts in physicochemical, metabolic, and OC character between vegetation states

Differences in vegetation states corresponded to differences in physicochemistry, aerobic metabolism, and OC pool composition. V was characterized by mature trees near the water line and was nutrient-rich relative to NV (Figure S1-3). V displayed comparatively high concentrations of total C and rates of aerobic metabolism (Figure S1-3). In contrast, NV consisted of vegetation-free, cobble-ridden shoreline with sandier soils, low total C, and low aerobic metabolism (Figure S1-3).

Compositional difference in OC pools indicated a possibility for distinct OC oxidation processes between the vegetation states (Figure 3, PERMANOVA, P = 0.001 in both water-soluble and bound-OC pools). If each pool contained similar OC compounds, one might infer similar OC inputs and biochemical processing in each vegetation state. Differences in OC pool composition within each vegetation state therefore putatively denotes either variation in OC inputs and/or variation in OC processing. Additionally, total OC content explained only 42% of aerobic metabolic rates across both sites (R^2^ = 0.42, P = 0.0009) and had no significant influence on metabolism within each site (V: P = 0.69, NV: P = 0.10, Figure S4). This suggests that factors beyond bulk OC content influenced aerobic metabolism. We hypothesized that variation in OC composition was responsible for variation in aerobic metabolism within each vegetation state. The following sections explore this possibility.

**Figure 3.**
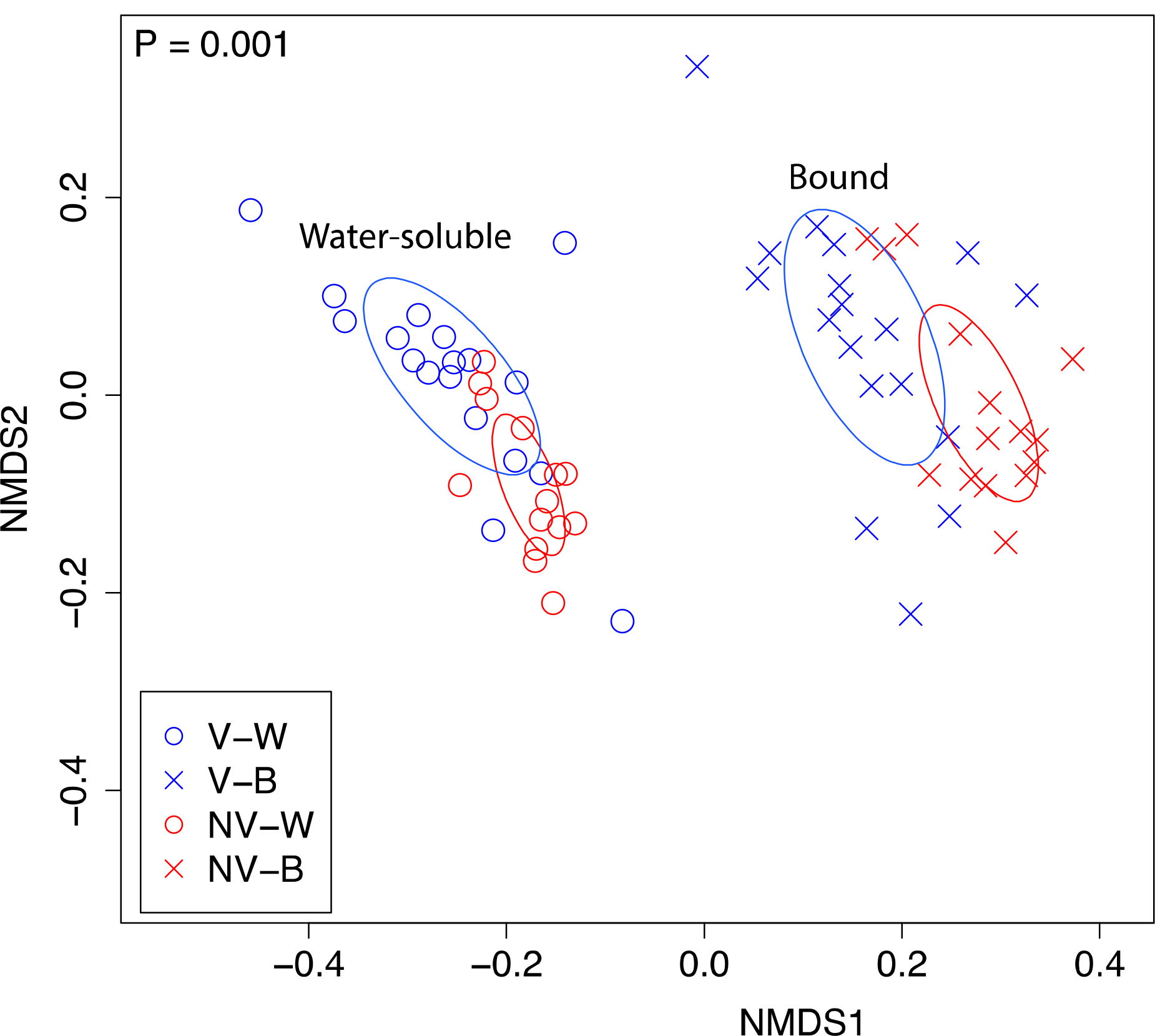
NMDS visualization of dissimilarity in OC pool composition. Water-soluble and bound-OC pools are represented by open circles and x’s, respectively. Samples associated with riparian vegetation are blue, and those in areas without vegetation are red. The P-value reflects differences among all groups, as assessed by PERMANOVA. Ellipses represent the standard deviation of the average axis scores for each group, generated using the ‘ordiellipse’ function in the ‘vegan’ package. Within each extraction, the composition of OC pools was significantly different across vegetation states (both P = 0.001).

### 3.2. Associations between C transformations and aerobic metabolism

Given compositional differences in OC between vegetation states and known impacts of C chemistry on metabolic functioning in other systems [*Castle et al.*, 2016; *Graham et al.*, 2016b], we hypothesized that biochemical transformations that were related to rates of aerobic metabolism would be unique to each vegetation state.

Consistent with this hypothesis, transformation analysis indicated that the biochemical processes associated with OC oxidation were significantly different between the vegetation states. Specifically, OC transformations correlated to aerobic metabolism were significantly different at each vegetation state (PERMANOVA, H_2_O P = 0.005 and CHCl_3_ P = 0.001, Figure 4 a-b, Table 2). In comparing differences in transformations occurring within the water-soluble OC pool, we observed higher abundances of amino- and sugar-associated transformations for V-W relative to NV-W. Twenty-six and four of these transformations co-varied significantly with aerobic metabolism in V-W and NV-W, respectively. These V-W transformations were primarily associated with simple C molecules (*e.g*., glucose, Table 2). Conversely, within the bound-OC pool, 35 transformations co-varied with aerobic metabolism in NV-B, compared to only three in V-B. In both cases, transformations associated with bound-OC consisted of a greater proportion of complex C molecules (*e.g*., biotinyl, palmitoylation, and glyoxylate, Table 2) than in water-soluble pools.

**Figure 4.**
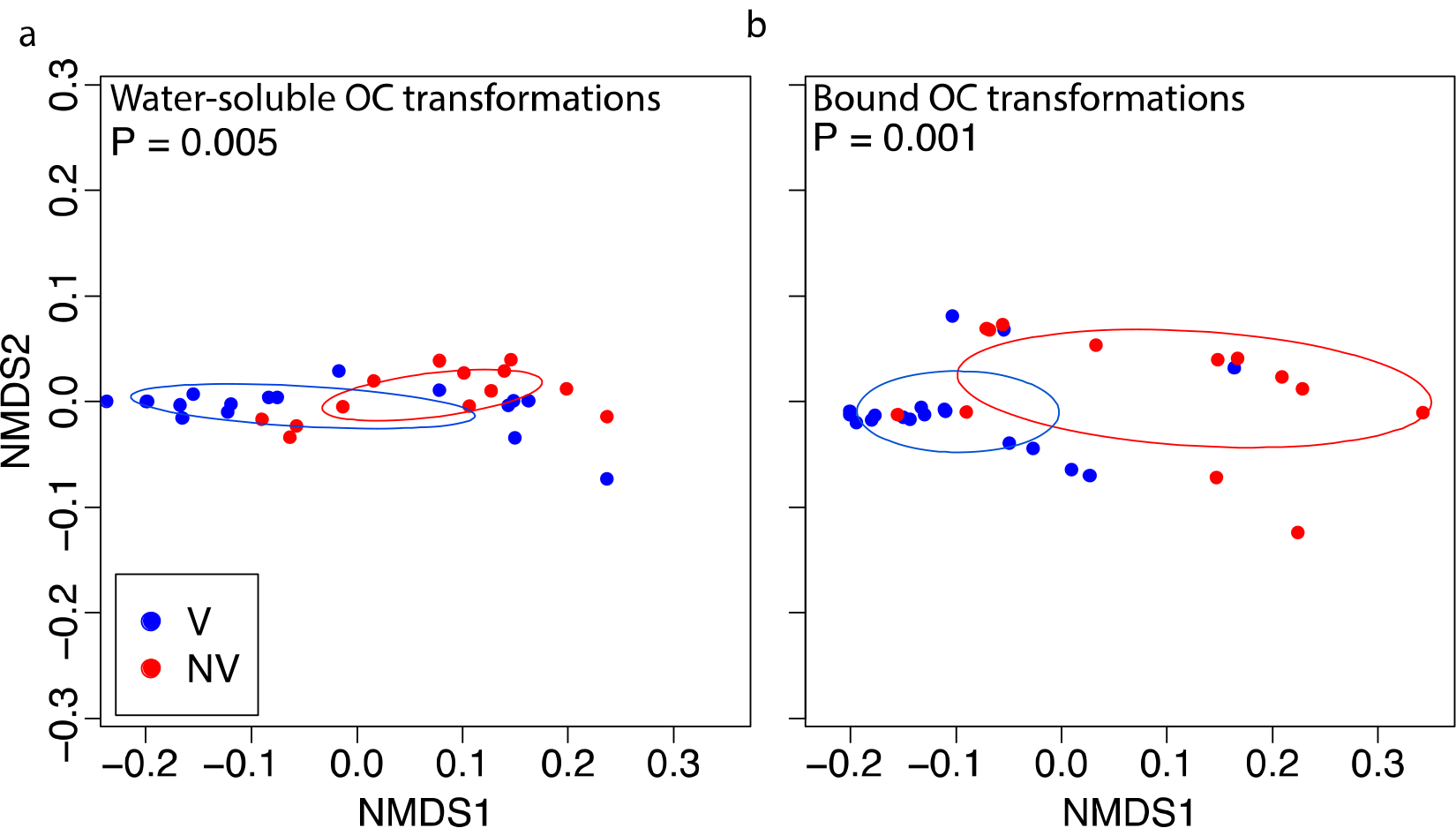
NMDS visualization of biochemical transformation partitioning among vegetation states. Biochemical transformations that were correlated to aerobic metabolism were significantly different among vegetation states in both the (a) water-soluble and (b) bound-OC pools. V and NV are denoted in blue and red, respectively, and significance values are derived from PERMANOVA. Ellipses represent the standard deviation of the average axis scores for each group, generated using the ‘ordiellipse’ function in the ‘vegan’ package.

**Table 2.** Biochemical transformations and the amount of variation 887 in aerobic metabolism explained by each (R^2^), within each OC pool and vegetation state.

| | $R^2$ |
| --- | --- |
| **V-W** |  |
| uridine_5_diphosphate_(-H2O)_C9H12N2O11P2 | 0.49 |
| cytosine_(-H)_C4H4N3O | 0.44 |
| uridine_5_monophosphate_(-H2O)_C9H11N2O8P | 0.41 |
| guanine_(-H)_C5H4N5O | 0.38 |
| guanosine_(-H2O)_C10H11N5O4 | 0.37 |
| adenine_(-H)_C5H4N5 | 0.36 |
| glutathione_(-H2O)_C10H15N3O5S | 0.34 |
| uracil_(-H)_C4H3N2O2 | 0.33 |
| C6H10O6 | 0.30 |
| glucose_C6H12O6 | 0.29 |
| Aspartic_Acid_C4H5NO3 | 0.28 |
| Glucuronic_Acid_(-H2O) | 0.28 |
| D-Ribose_(-H2O)_(-ribosylation) | 0.27 |
| secondary_amine | 0.26 |
| C6H10O5 | 0.25 |
| monosaccharide_(-H2O) | 0.25 |
| Glutamic_Acid_C5H7NO3 | 0.24 |
| diphosphate_H3O6P2 | 0.23 |
| thymine_(-H)_C5H5N2O2 | 0.23 |
| glyoxylate_(-H2O)_C2O2 | 0.23 |
| malonyl_group_(-H2O)_C3H2O3 | 0.22 |
| adenosine_(-H2O)_C10H11N5O3 | 0.22 |
| primary_amine | 0.21 |
| thymidine_(-H2O)_C10H12N2O4 | 0.21 |
| carbamoyl_P_transfer_(-H2PO4)_CH2ON | 0.20 |
| erythrose_(-H2O) | 0.20 |
| **V-B** |  |
| isoprene_addition_(-H)_C5H7 | 0.39 |
| hydrogenation_dehydrogenation_H2 | 0.24 |
| biotinyl_(-H2O)_C10H14N2O2S | 0.16 |
| **NV-W** |  |
| Glucuronic_Acid_(-H2O) | 0.49 |
| C6H10O6 | 0.49 |
| acetone_(-H)_C3H5O | 0.45 |
| adenine_(-H)_C5H4N5 | 0.45 |

The larger number of transformations associated with aerobic metabolism in V-W vs. V-B, and the larger number in NV-B vs. NV-W, suggests that aerobic metabolism in vegetated and unvegetated areas depend on water-soluble and bound-OC pools, respectively. We note some oxidation of the bound-OC pool under vegetated conditions, but only three correlations were observed between V-B transformations and aerobic metabolism, suggesting a relatively minor role, especially considering that there were 35 significant correlations for NV-B.

These differences suggest that an increased supply of bioavailable compounds in vegetated areas leads to bound-OC being less involved in aerobic metabolism, relative to unvegetated areas where bound-OC appears to be heavily involved in aerobic metabolism. The concept of priming [*Kuzyakov*, 2010] would predict the opposite pattern; a greater supply of bioavailable OC should increase the contributions of less bioavailable OC (here, bound-OC) to aerobic metabolism. Our results run counter to a priming mechanism and indicate that the supply of bioavailable compounds diminishes the contribution of bound-OC to aerobic metabolism and, in turn, protects bound-OC pools. Conversely, metabolism of bound-OC pools in locations without riparian vegetation possibly denotes microbial adaptations to a low-C environment that supports the oxidation of mineral-bound OC. Mineral-bound OC therefore has greater potential to remain sequestered along river corridors with spatially and temporally consistent inputs of bioavailable OC, potentially derived from riparian vegetation.

### 3.3. Associations between microbial metabolic pathways and aerobic metabolism

Because we observed stark differences in the identity of OC transformations that correlated with aerobic metabolism across vegetation states, we hypothesized that the microbial metabolic pathways associated with OC transformations were also dependent on vegetation state. Indeed, pathways associated with OC oxidation were distinct at V vs. NV, supporting our hypothesis that there were differences in the metabolic processing of OC in the presence or absence of riparian vegetation. Specifically, while the metabolism of plant-derived compounds appeared to be a major driver of aerobic respiration at both vegetation states, metabolism at V mostly involved readily bioavailable plant derivatives in the water-soluble OC pool, and metabolism at NV was associated with plant derivatives in the bound-OC pool (Figure 5).

**Figure 5.**
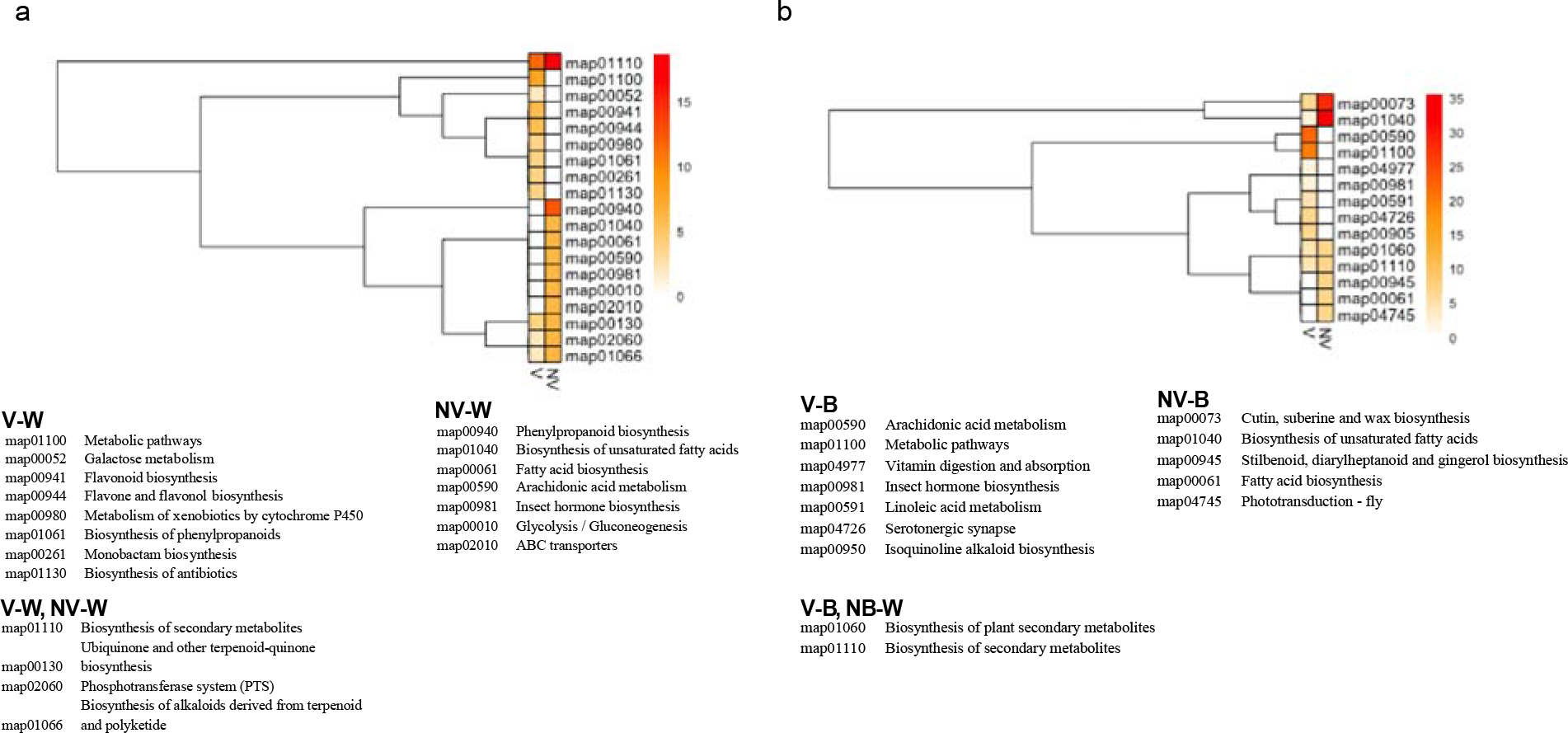
KEGG pathways associated with aerobic metabolism. A hierarchical clustering heatmap shows KEGG pathways associated with aerobic metabolism. (a) and (b) show water-soluble and bound-OC pools, respectively. Colors move from white to red on a scale of 0% to 20% in (a) and 0% to 35% in (b), showing percent relative abundance of each pathway in each group. Pathways are described in the legends. Pathways listed under each condition (e.g., V-W, etc.) had significant correlations to aerobic metabolism within that condition.

In V-W, pathways that corresponded to aerobic metabolism included map01110 biosynthesis of secondary metabolites, map000941 flavonoid biosynthesis, and map00944 flavone and flavonol biosynthesis, among others (see Figure 5a). Each of these pathways denotes an association between plant-derived compounds and OC oxidation in sediments. Secondary metabolites (map01110) are largely comprised of plant-derived compounds such as flavonoids [*Agati et al.*, 2012], terpenoids [*Tholl*, 2015], and nitrogen-containing alkaloids [*Willaman and Schubert*, 1961], while flavonoids [*Agati et al.*, 2012] are one of the most abundant plant-derived compounds. Associations with flavone/flavonol [*Agati et al.*, 2012] and phenylpropanoids [*Hahlbrock and Scheel*, 1989] bolster a possible relationship between plant-associated metabolites and aerobic metabolism in V-W.

Although correlations between plant-associated KEGG pathways and aerobic metabolism could indicate the persistence of plant secondary metabolites rather than microbial metabolism, our results indicate a central role for vegetation in water-soluble OC oxidation in either case. For example, if KEGG associations were attributable to plant metabolism instead of microbial metabolism, correlations between plant-associated pathways and aerobic metabolism in V-W would indicate an indirect relationship between plant growth and microbial oxidation of OC, whereby plant byproducts support microbial communities in oxidizing other portions of the OC pool.

Pathways correlated with aerobic metabolism in NV-W or V-W had some overlap, but were mostly distinct from each other (Figure 5a). Although some correlations in NV-W indicated associations between plant-associated metabolic pathways and OC oxidation (*e.g*., map01110), relationships also indicated the involvement of broad metabolic processes including membrane transport (map02010 ABC transporters), carbohydrate metabolism (map00010 Glycolysis/Gluconeogensis), and lipid metabolism (map00061 Fatty acid biosynthesis and map01040 Biosynthesis of unsaturated fatty acids). We propose that these associations indicate utilization of alternative resources in comparision to V-W, in which relationships almost exclusively denoted the metabolism of readily bioavailable plant byproducts. In particular, we note correlations in NV-W between OC oxidation and lipid-based molecules (map00061 and map01040) that are typically poorly-represented in water-soluble OC pools.

We also observed largely distinct relationships between metabolisms and OC oxidation in V-B versus NV-B. The lipid-based and plant-associated pathway map00073 (Cutin, suberine, and wax biosynthesis [*King et al.*, 2007; *Raffaele et al.*, 2009; *Shepherd and Wynne Griffiths*, 2006]) comprised almost 35% of relationships in NV-B, and more generic plant-associated pathways such as map01060 (Biosynthesis of plant secondary metabolites) and map01110 (Biosynthesis of secondary metabolites) were also prevalent (Figure 5b). Conversely in V-B, the most abundant associations were non-lipid metabolisms map00590 (Arachidonic acid metabolism) and map01100 (Metabolic pathways).

Together, our results indicate largely separate metabolic processes occurring in each of the water-soluble and bound-OC pools in V and NV. Because of the abundance of lipid-based metabolisms correlated to OC oxidation at NV (in both OC pools), we further hypothesize that these metabolisms comprise the primary KEGG-identifiable pathways associated with OC oxidation in areas without riparian vegetation. This contrasts to the cycling of a more diverse range of secondary metabolite classes in areas with more significant OC inputs from riparian vegetation. We note that correlations between KEGG pathways and aerobic metabolism in V-B indicates less oxidation of lipid plant material in the bound-OC pool under vegetated conditions than in NV-B. We therefore propose that plant-derived lipid compounds serve as a secondary substrate for OC oxidation in shorelines with riparian vegetation, given that more correlations at V were detected in the water-soluble pool.

### 3.4. Thermodynamics of carbon oxidation

Finally, we hypothesized that microbes would preferentially oxidize more thermodynamically favorable compounds at both sites, consistent with common thermodynamic constraints on biogeochemical cycles [*Burgin et al.*, 2011; *Hedin et al.*, 1998; *Helton et al.*,2015]. Because we observed evidence for preferential OC oxidation of the water-soluble OC pool at V and of the bound-OC pool at NV, we further hypothesized that thermodynamic-based preference of OC oxidation would be observable only in the preferred substrate pool within each vegetation state. Consistent with this hypothesis, aerobic metabolism was positively correlated to average 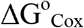 in V-W (R^2^ = 0.37, P = 0.008, Figure 6a) and NV-B (R^2^ = 0.59, P = 0.0005 Figure 6b), but these variables were not correlated in V-B or NV-W. In both cases, aerobic metabolism corresponded to a depletion of more thermodynamically favorable OC (*i.e*., OC became less favorable as aerobic metabolism increased), resulting in progressively less favorable thermodynamic conditions.

**Figure 6.**
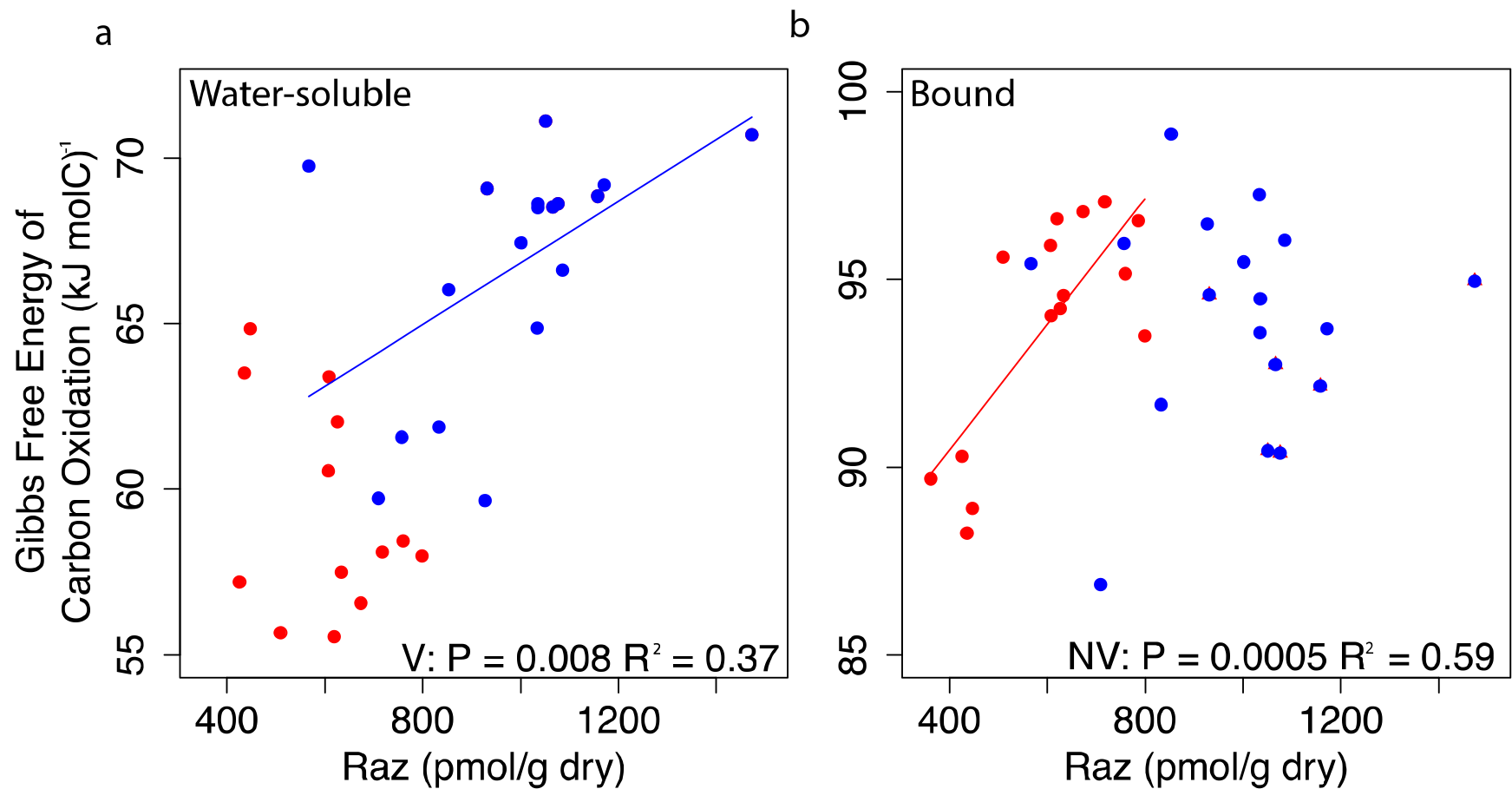
Correlations between Gibbs free energy of carbon oxidation (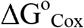) and aerobic metabolism. (a) and (b) display mixed linear regressions relating 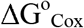 to aerobic metabolism in water-soluble and bound-OC pools, respectively. Aerobic metabolism is expressed as pmoles of resazurin reduced to resorufin per gram of sediment dry weight over a 48hr incubation period (Raz, see Supplemental Information). V and NV are denoted in blue and red, and solid lines show significant relationships within each vegetation state.

The priming conceptual framework would predict that terrestrial inputs associated with riparian vegetation should condition microbial communities to oxidize less thermodynamically favorable C, such as that found in the bound-OC pool. In such a scenario, inputs of thermodynamically favorable carbon should—by minimizing community-level energy constraints—allow for the rise of microbial physiologies that can oxidize less favorable C [*Kuzyakov*, 2010]. In this case, a significant relationship between thermodynamic favorability and aerobic metabolism in the V-W pool should lead to a similar relationship within the V-B pool. Our results reveal a strong relationship within the V-W pool, but not in the V-B pool, thereby rejecting an influence of priming. Instead, our results suggest that bound-OC pools are protected by thermodynamically favorable compounds that serve as preferred substrate.

In contrast to our expectation that water-soluble OC associated with riparian vegetation would increase oxidation of bound-OC pools, we observed evidence consistent with inhibition of bound-OC oxidation by thermodynamically favorable water-soluble compounds. Priming has been actively debated in aquatic research [*Bengtsson et al.*, 2014; *Bianchi*, 2011; *Guenet et al.*, 2010], and a number of other studies have been unable to detect a priming effect in sediment and aqueous habitats [*Bengtsson et al.*, 2014; *Catalán et al*, 2015].

### 3.5. Conceptual model for OC oxidation at terrestrial-aquatic interfaces

Based on our work, we propose a new view of OC oxidation along terrestrial-aquatic interfaces in which the oxidation of bound-OC is limited by terrestrial inputs from riparian vegetation (Figure 7). While our work represents a single system, our conceptual model is derived from ultra-high resolution measurements that provide greater mechanistic insight into OC cycles than traditional measures. Our model is therefore meant to provide a basis for more spatially extensive experiments that will allow for broader transferability.

**Figure 7.**
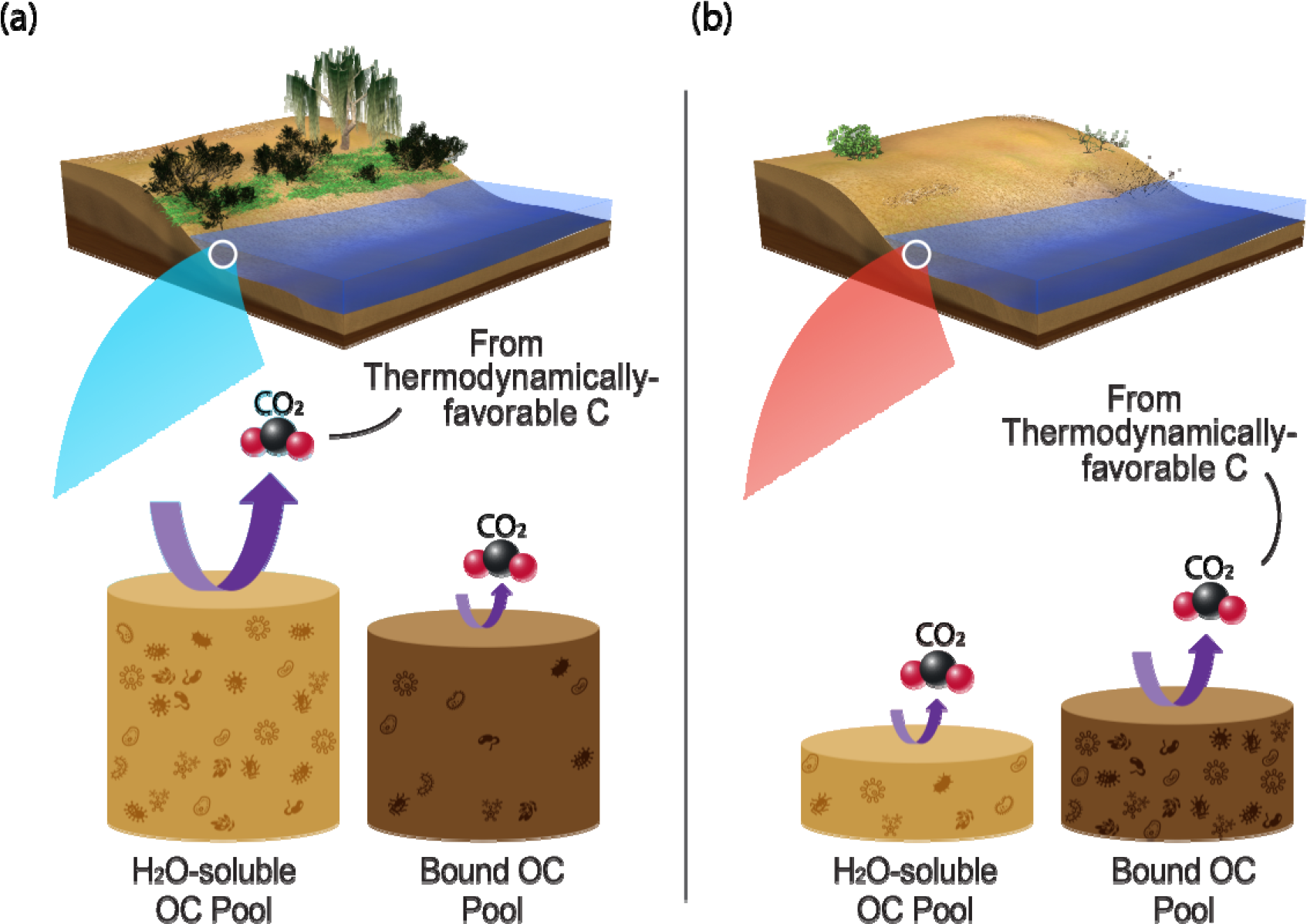
Conceptualization of relationship between riparian vegetation and OC oxidation. We propose a conceptualization of OC oxidation at terrestrial-aquatic interfaces whereby (a) more riparian vegetation results in greater terrestrial C deposition and larger water-soluble and bound-OC pools. However, water-soluble OC is preferentially oxidized, which protects the bound-OC pool. Conversely, (b) areas deplete in riparian vegetation experience lower inputs of water-soluble OC and show lower rates of OC oxidation. This results in smaller OC pools (water-soluble and bound) and microbial adaptation for oxidation of the bound-OC pool. In both cases, the most thermodynamically favorable portions of the OC pool being metabolized are preferentially oxidized. Height of the cylinders denotes pool sizes, and arrow thickness denotes flux magnitude.

We hypothesize that riparian vegetation sustains inputs of water-soluble compounds to nearshore OC pools, resulting in a larger thermodynamically favorable, water-soluble OC pool (Figure 7b). This leads to higher overall C content in nearshore sediments and elevated rates of aerobic respiration relative to areas with less riparian vegetation. Our data also suggest that in the presence of riparian vegetation, microbial carbon oxidation primarily uses the water-soluble OC pool with minimal oxidation of bound-OC due to physical and/or thermodynamic protection of this pool. For instance, 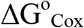 was lower in water-soluble OC pools than in bound-OC, and a large presence of this thermodynamically favorable pool may provide adequate substrate to sustain metabolic functioning, limiting the need to metabolize less thermodynamically favorable OC. Additionally, organomineral interactions can protect bound-OC from extracellular enzyme acitivity [*Hunter et al.*, 2016], inhibiting the bioavailability of OC.

In contrast, non-vegetated riparian zones provide little input into water-soluble OC pools (Figure 7a), and rates of metabolism and C pool sizes are lower in these environments. Carbon oxidation in these non-vegetated zones occurs primarily within the bound-OC pool, albeit more slowly and as product of different biochemical and metabolic pathways than in vegetated environments (*e.g*., complex C transformations and lipid-based metabolism of plant derivatives at NV). We posit that water-soluble pools in non-vegetated sediments are sufficiently small that investing in enzymes needed to metabolize this OC pool results in a net energy loss. Instead, microbes in unvegetated areas must invest in cellular machinery to access bound-OC, and our results imply that the cellular machinery needed to access bound-OC is distinct from the machinery needed to access water-soluble OC.

Interestingly, aerobic metabolism within both types of sediments is related to a depletion of thermodynamically favorable compounds; however, this occurs in water-soluble OC pools in vegetated zones and bound-OC pools in non-vegetated zones. That is, microorganisms in both environments are constrained to the metabolism of their primary substrate pool but preferentially oxidize more thermodynamically favorable compounds within that pool. This suggests that microorganisms are conditioned to metabolize a subset of compounds within sediment OC, possibly defined by thermodynamic or physical protection mechanisms, but operate under common thermodynamic constraints once adapted to oxidize a certain OC pool.

### 3.6. Broader Implications

Our work is particularly relevant to CO_2_ emissions in light of changes in land cover and precipitation, and the influence of these perturbations to on C fluxes across terrestrial-aquatic interfaces. Carbon moving across terrestrial-aquatic interfaces has been examined primarily for its propensity to be oxidized along land-to-sea continuums [*Battin et al.*, 2008; *Battin et al.*, 2009; *Regnier et al.*, 2013], but we suggest that this terrestrially-derived OC also has important influences on stabilizing mineral-bound OC within nearshore environments. The magnitude, distribution, and chemical quality of terrestrial C fluxes into aquatic environments are perturbed by shifts in land cover [*Fang et al.*, 2005; *Knapp et al.*, 2008]. Furthermore, vegetation distributions in natural ecosystems are predicted to shift in response to altered precipitation regimes. Associated changes in plant phenology, morphology, and establishment will impact the quantity, quality, and distribution of terrestrial material entering aquatic systems [*Knapp et al.*, 2008], and we currently have an incomplete understanding of how these patterns will vary across ecosystems and precipitation patterns [*Fang et al.*, 2005; *Knapp et al.*, 2008]. A mechanistic framework for C oxidation that captures impacts of heterogeneity in vegetation in river corridors will therefore aid in predicting how terrestrial-aquatic interfaces respond to ongoing perturbations. Here, we demonstrate that increases in the flux of terrestrial C to aquatic systems may lead to larger mineral-bound C pools by physically and thermodynamically protecting these pools. Conversely, we demonstrate a potential for oxidation of mineral-bound C pools in areas with diminished terrestrial C inputs.

Earth System Models (ESMs) depend on mathematical representations of C cycling, and the continued development of these models is tightly coupled to conceptual advances drawn from field-based observations [*Burd et al.*, 2016; *Six et al.*, 2002]. Despite recent progress, these models are still missing key regulatory processes [*Todd-Brown et al.*, 2013; *Wieder et al.*, 2013; *Wieder et al.*, 2015]. In particular, terrestrial-aquatic interfaces are now considered to be active zones of OC transformation as opposed to their representation as non-reactive ‘pipes’ in most ESMs. To address this knowledge gap, we propose a new conceptual framework of OC dynamics based on analysis of *in situ* observational data that explicitly considers a central challenge in model improvement—biochemical, metabolic, and thermodynamic mechanisms governing OC oxidation along terrestrial-aquatic interfaces. Our results directly contrast those expected within a ‘priming’ framework, and we advance that water-soluble thermodynamically favorable OC associated with riparian vegetation protects thermodynamically less favorable bound-OC from oxidation. We also demonstrate differences in biochemical and metabolic pathways associated with metabolism of water-soluble and bound-OC pools in the presence or absence of riparian vegetation, furthering a processed-based understanding of terrestrial-aquatic interfaces. Our research provides an opportunity to enhance the mechanistic underpinning of OC oxidation process representations within ESMs and proposes interactions between OC thermodynamics and mineral-inhibition of OC oxidation as a key future research need.

## Author Contributions

EBG was responsible for conceptual development and data analysis and was the primary writer with guidance from JCS and MT. ARC, AEG, CTR, ECR, DWK, and JCS were responsible for experimental design and data collection. MT was responsible for all FTICR processing, and LB assisted with data analysis. All authors contributed to manuscript revisions.

## Acknowledgements

This research was supported by the US Department of Energy (DOE), Office of Biological and Environmental Research (BER), as part of Subsurface Biogeochemical Research Program’s Scientific Focus Area (SFA) at Pacific Northwest National Laboratory (PNNL). PNNL is operated for DOE by Battelle under contract DE-AC06-76RLO 1830. A portion of the research was performed at the Environmental Molecular Science Laboratory User Facility located on PNNL’s campus. We thank Nancy Hess for helpful feedback in manuscript revision and Jeff Holmes for assistance in text editing. All data are publicly available at DOI XXXXXXX.

## References

Aalto, R., L. Maurice-Bourgoin, T. Dunne, D. R. Montgomery, C. A. Nittrouer, and J.-L. Guyot (2003), Episodic sediment accumulation on Amazonian flood plains influenced by El Nino/Southern Oscillation, Nature, 425(6957), 493–497.

Agati, G., E. Azzarello, S. Pollastri, and M. Tattini (2012), Flavonoids as antioxidants in plants: location and functional significance, Plant Science, 196, 67–76.

Arndt, S., B. B. Jørgensen, D. E. LaRowe, J. Middelburg, R. Pancost, and P. Regnier (2013), Quantifying the degradation of organic matter in marine sediments: a review and synthesis, Earth-science reviews, 123, 53–86.

Arntzen, E. V., D. R. Geist, and P. E. Dresel (2006), Effects of fluctuating river flow on groundwater/surface water mixing in the hyporheic zone of a regulated, large cobble bed river, River Research and Applications, 22(8), 937–946.

Bailey, V. L., A. Smith, M. Tfaily, S. J. Fansler, and B. Bond-Lamberty (2017), Differences in soluble organic carbon chemistry in pore waters sampled from different pore size domains, Soil Biology and Biochemistry, 107, 133–143.

Battin, T. J., L. A. Kaplan, S. Findlay, C. S. Hopkinson, E. Marti, A. I. Packman, J. D. Newbold, and F. Sabater (2008), Biophysical controls on organic carbon fluxes in fluvial networks, Nature Geoscience, 1(2), 95–100.

Battin, T. J., S. Luyssaert, L. A. Kaplan, A. K. Aufdenkampe, A. Richter, and L. J. Tranvik (2009), The boundless carbon cycle, Nature Geoscience, 2(9), 598–600.

Bengtsson, M. M., K. Wagner, N. R. Burns, E. R. Herberg, W. Wanek, L. A. Kaplan, and T. J. Battin (2014), No evidence of aquatic priming effects in hyporheic zone microcosms, Scientific reports, 4, 5187.

Bianchi, T. S. (2011), The role of terrestrially derived organic carbon in the coastal ocean: A changing paradigm and the priming effect, Proceedings of the National Academy of Sciences, 108(49), 19473–19481.

Blagodatskaya, E., and Y. Kuzyakov (2008), Mechanisms of real and apparent priming effects and their dependence on soil microbial biomass and community structure: critical review, Biology and Fertility of Soils, 45(2), 115–131.

Breitling, R., S. Ritchie, D. Goodenowe, M. L. Stewart, and M. P. Barrett (2006), Ab initio prediction of metabolic networks using Fourier transform mass spectrometry data, Metabolomics, 2(3), 155–164.

Burd, A. B., S. Frey, A. Cabre, T. Ito, N. M. Levine, C. Lønborg, M. Long, M. Mauritz, R. Q. Thomas, and B. M. Stephens (2016), Terrestrial and marine perspectives on modeling organic matter degradation pathways, Global change biology, 22(1), 121–136.

Burgin, A. J., W. H. Yang, S. K. Hamilton, and W. L. Silver (2011), Beyond carbon and nitrogen: how the microbial energy economy couples elemental cycles in diverse ecosystems, Frontiers in Ecology and the Environment, 9(1), 44–52.

Castle, S. C., D. R. Nemergut, A. S. Grandy, J. W. Leff, E. B. Graham, E. Hood, S. K. Schmidt, K. Wickings, and C. C. Cleveland (2016), Biogeochemical drivers of microbial community convergence across actively retreating glaciers, Soil Biology and Biochemistry, 101, 74–84.

Catalán, N., A. M. Kellerman, H. Peter, F. Carmona, and L. J. Tranvik (2015), Absence of a priming effect on dissolved organic carbon degradation in lake water, Limnology and Oceanography, 60(1), 159–168.

Cotrufo, M. F., M.D. Wallenstein, C. M. Boot, K. Denef, and E. Paul (2013), The Microbial Efficiency - Matrix Stabilization (MEMS) framework integrates plant litter decomposition with soil organic matter stabilization: do labile plant inputs form stable soil organic matter?, Global Change Biology, 19(4), 988–995.

Dorado-García, I., J. Syväranta, S. P. Devlin, J. M. Medina-Sánchez, and R. I. Jones (2016), Experimental assessment of a possible microbial priming effect in a humic boreal lake, Aquatic Sciences, 78(1), 191–202.

Ebel, W. J., C. D. Becker, J. W. Mullan, and H. L. Raymond (1989), The Columbia River–toward a holistic understanding, Canadian special publication of fisheries and aquatic sciences/Publication speciale canadienne des sciences halieutiques et aquatiques, 1989.

Fang, J., S. Piao, L. Zhou, J. He, F. Wei, R. B. Myneni, C. J. Tucker, and K. Tan (2005), Precipitation patterns alter growth of temperate vegetation, Geophysical research letters, 32(21).

Graham, E. B., A. R. Crump, C. T. Resch, S. Fansler, E. Arntzen, D. W. Kennedy, J. K. Fredrickson, and J. C. Stegen (2016a), Coupling spatiotemporal community assembly processes to changes in microbial metabolism, Frontiers in Microbiology, 7, 1949.

Graham, E. B., A. R. Crump, C. T. Resch, S. Fansler, E. Arntzen, D. W. Kennedy,J. K. Fredrickson, and J. C. Stegen (2017), Deterministic influences exceed dispersal effects on hydrologically - connected microbiomes, Environmental Microbiology, 19(4), 1552–1567.

Graham, E. B., J. E. Knelman, R. S. Gabor, S. Schooler, D. M. McKnight, and D. Nemergut (2016b), Dissolved organic matter and inorganic mercury loadings favor novel methylators and fermentation metabolisms in oligotrophic sediments, bioRxiv, 072017.

Guenet, B., M. Danger, L. Abbadie, and G. Lacroix (2010), Priming effect: bridging the gap between terrestrial and aquatic ecology, Ecology, 91(10), 2850–2861.

Haggerty, R., E. Martí, A. Argerich, D. Von Schiller, and N. B. Grimm (2009), Resazurin as a “smart” tracer for quantifying metabolically active transient storage in stream ecosystems, Journal of Geophysical Research: Biogeosciences, 114(G3).

Hahlbrock, K., and D. Scheel (1989), Physiology and molecular biology of phenylpropanoid metabolism, Annual review of plant biology, 40(1), 347–369.

Hedges, J., and J. Oades (1997), Comparative organic geochemistries of soils and marine sediments, Organic geochemistry, 27(7), 319–361.

Hedges, J. I., G. Eglinton, P. G. Hatcher, D. L. Kirchman, C. Arnosti, S. Derenne, R. P. Evershed, I. Kögel-Knabner, J. De Leeuw, and R. Littke (2000), The molecularly-uncharacterized component of nonliving organic matter in natural environments, Organic Geochemistry, 31(10), 945–958.

Hedges, J. I., and R. G. Keil (1995), Sedimentary organic matter preservation: an assessment and speculative synthesis, Marine chemistry, 49(2-3), 81–115.

Hedin, L. O., J. C. von Fischer, N. E. Ostrom, B. P. Kennedy, M. G. Brown, and G. P. Robertson (1998), Thermodynamic constraints on nitrogentransformations and other biogeochemical processes at soil–stream interfaces, Ecology, 79(2), 684–703.

Helton, A. M., M. Ardòn, and E. S. Bernhardt (2015), Thermodynamic constraints on the utility of ecological stoichiometry for explaining global biogeochemical patterns, Ecology letters, 18(10), 1049–1056.

Herzsprung, P., K. Osterloh, W. von Tümpling, M. Harir, N. Hertkorn, P. Schmitt-Kopplin, R. Meissner, S. Bernsdorf, and K. Friese (2017), Differences in DOM of rewetted and natural peatlands–Results from high-field FT-IFTICR-MS and bulk optical parameters, Science of The Total Environment, 586, 770–781.

Hunter, W. R., R. Niederdorfer, A. Gernand, B. Veuger, J. Prommer, M. Mooshammer, W. Wanek, and T. J. Battin (2016), Metabolism of mineral - sorbed organic matter and microbial lifestyles in fluvial ecosystems, Geophysical Research Letters.

Kanehisa, M., and S. Goto (2000), KEGG: kyoto encyclopedia of genes and genomes, Nucleic acids research, 28(1), 27–30.

Kellerman, A. M., D. N. Kothawala, T. Dittmar, and L. J. Tranvik (2015), Persistence of dissolved organic matter in lakes related to its molecular characteristics, Nature Geoscience, 8(6), 454–457.

Kim, S., R. W. Kramer, and P. G. Hatcher (2003), Graphical method for analysis of ultrahigh-resolution broadband mass spectra of natural organic matter, the van Krevelen diagram, Analytical Chemistry, 75(20), 5336–5344.

King, A., J.-W. Nam, J. Han, J. Hilliard, and J. G. Jaworski (2007), Cuticular wax biosynthesis in petunia petals: cloning and characterization of an alcohol-acyltransferase that synthesizes wax-esters, Planta, 226(2), 381–394.

Knapp, A. K., C. Beier, D. D. Briske, A. T. Classen, Y. Luo, M. Reichstein, M. D. Smith, S. D. Smith, J. E. Bell, and P. A. Fay (2008), Consequences of more extreme precipitation regimes for terrestrial ecosystems, Bioscience, 58(9), 811–821.

Koch, B. P., M. Witt, R. Engbrodt, T. Dittmar, and G. Kattner (2005), Molecular formulae of marine and terrigenous dissolved organic matter detected by electrospray ionization Fourier transform ion cyclotron resonance mass spectrometry, Geochimica et Cosmochimica Acta, 69(13), 3299–3308.

Koégel-Knabner, I. (2002), The macromolecular organic composition of plant and microbial residues as inputs to soil organic matter, Soil Biology and Biochemistry, 34(2), 139–162.

Kögel-Knabner, I. (2000), Analytical approaches for characterizing soil organic matter, Organic Geochemistry, 31(7), 609–625.

Kujawinski, E. B., and M. D. Behn (2006), Automated analysis of electrospray ionization Fourier transform ion cyclotron resonance mass spectra of natural organic matter, Analytical Chemistry, 78(13), 4363–4373.

Kuzyakov, Y. (2010), Priming effects: interactions between living and dead organic matter, Soil Biology and Biochemistry, 42(9), 1363–1371.

LaRowe, D. E., and P. Van Cappellen (2011), Degradation of natural organic matter: a thermodynamic analysis, Geochimica et Cosmochimica Acta, 75(8), 2030–2042.

Lehmann, J., and M. Kleber (2015), The contentious nature of soil organic matter, Nature, 528(7580), 60–68.

Lin, X., J. McKinley, C. T. Resch, R. Kaluzny, C. L. Lauber, J. Fredrickson, R. Knight, and A. Konopka (2012), Spatial and temporal dynamics of the microbial community in the Hanford unconfined aquifer, The ISME Journal, 6(9), 1665–1676.

Lotspeich, F. B., and B. H. Reid (1980), Tri-tube freeze-core procedure for sampling stream gravels, The Progressive Fish-Culturist, 42(2), 96–99.

Mason, H., J. Begg, R. S. Maxwell, A. B. Kersting, and M. Zavarin (2016), A novel solid-state NMR method for the investigation of trivalent lanthanide sorption on amorphous silica at low surface loadings, Environmental Science: Processes & Impacts.

Minor, E. C., C. J. Steinbring, K. Longnecker, and E. B. Kujawinski (2012), Characterization of dissolved organic matter in Lake Superior and its watershed using ultrahigh resolution mass spectrometry, Organic Geochemistry, 43, 1–11.

Moser, D. P., J. K. Fredrickson, D. R. Geist, E. V. Arntzen, A. D. Peacock, S.-M. W. Li, T. Spadoni, and J. P. McKinley (2003), Biogeochemical processes and microbial characteristics across groundwater-surface water boundaries of the Hanford Reach of the Columbia River, Environmental Science & Technology, 37(22), 5127–5134.

Peterson, R. E., and M. P. Connelly (2004), Water movement in the zone of interaction between groundwater and the Columbia River, Hanford site, Washington, Journal of Hydraulic Research, 42(S1), 53–58.

Raffaele, S., A. Leger, and D. Roby (2009), Very long chain fatty acid and lipid signaling in the response of plants to pathogens, Plant signaling & behavior, 4(2), 94–99.

Regnier, P., P. Friedlingstein, P. Ciais, F. T. Mackenzie, N. Gruber, I. A. Janssens, G. G. Laruelle, R. Lauerwald, S. Luyssaert, and A. J. Andersson (2013), Anthropogenic perturbation of the carbon fluxes from land to ocean, Nature geoscience, 6(8), 597–607.

Rood, K., and M. Church (1994), Modified freeze-core technique for sampling the permanently wetted streambed, North American Journal of Fisheries Management, 14(4), 852–861.

Rossel, P. E., C. Bienhold, A. Boetius, and T. Dittmar (2016), Dissolved organic matter in pore water of Arctic Ocean sediments: Environmental influence on molecular composition, Organic Geochemistry, 97, 41–52.

Rothman, D. H., and D. C. Forney (2007), Physical model for the decay and preservation of marine organic carbon, Science, 316(5829), 1325–1328.

Schmidt, M. W., M. S. Torn, S. Abiven, T. Dittmar, G. Guggenberger, I. A. Janssens, M. Kleber, I. Kögel-Knabner, J. Lehmann, and D. A. Manning (2011), Persistence of soil organic matter as an ecosystem property, Nature, 478(7367), 49–56.

Shepherd, T., and D. Wynne Griffiths (2006), The effects of stress on plant cuticular waxes, New Phytologist, 171(3), 469–499.

Six, J., P. Callewaert, S. Lenders, S. De Gryze, S. Morris, E. Gregorich, E. Paul, and K. Paustian (2002), Measuring and understanding carbon storage in afforested soils by physical fractionation, Soil science society of America journal, 66(6), 1981–1987.

Slater, L. D., D. Ntarlagiannis, F. D. Day-Lewis, K. Mwakanyamale, R. J. Versteeg, A. Ward, C. Strickland, C. D. Johnson, and J. W. Lane (2010), Use of electrical imaging and distributed temperature sensing methods to characterize surface water–groundwater exchange regulating uranium transport at the Hanford 300 Area, Washington, Water Resources Research, 46(10).

Stegen, J. C., J. K. Fredrickson, M. J. Wilkins, A. E. Konopka, W. C. Nelson, E. V. Arntzen, W. B. Chrisler, R. K. Chu, R. E. Danczak, and S. J. Fansler (2016), Groundwater-surface water mixing shifts ecological assembly processes and stimulates organic carbon turnover, Nature Communications, 7.

Stegen, J. C., X. Lin, A. E. Konopka, and J. K. Fredrickson (2012), Stochastic and deterministic assembly processes in subsurface microbial communities, The ISME Journal, 6(9), 1653–1664.

Tfaily, M. M., R. K. Chu, N. Tolić, K. M. Roscioli, C. R. Anderton, L. Pas⏑a-Tolić, E. W. Robinson, and N. J. Hess (2015), Advanced solvent based methods for molecular characterization of soil organic matter by high-resolution mass spectrometry, Analytical chemistry, 87(10), 5206–5215.

Tfaily, M. M., D. C. Podgorski, J. E. Corbett, J. P. Chanton, and W. T. Cooper (2011), Influence of acidification on the optical properties and molecular composition of dissolved organic matter, Analytica chimica acta, 706(2), 261–267.

Tfaily, M. M., P. Reardon, R. K. Chu, N. Tolić, L. Pas⏑a-Tolić, E. W. Robinson, and N. J. Hess (2017), Sequential extraction protocol for organic matter from soils and sediments using high resolution mass spectrometry and proton NMR, Analytica Chimica Acta.

Tholl, D. (2015), Biosynthesis and biological functions of terpenoids in plants, in Biotechnology of Isoprenoids, edited, pp. 63–106, Springer.

Todd-Brown, K., J. Randerson, W. Post, F. Hoffman, C. Tarnocai, E. Schuur, and S. Allison (2013), Causes of variation in soil carbon simulations from CMIP5 Earth system models and comparison with observations, Biogeosciences, 10(3).

Tremblay, L. B., T. Dittmar, A. G. Marshall, W. J. Cooper, and W. T. Cooper (2007), Molecular characterization of dissolved organic matter in a North Brazilian mangrove porewater and mangrove-fringed estuaries by ultrahigh resolution Fourier transform-ion cyclotron resonance mass spectrometry and excitation/emission spectroscopy, Marine chemistry, 105(1), 15–29.

Ward, C. P., and R. M. Cory (2015), Chemical composition of dissolved organic matter draining permafrost soils, Geochimica et Cosmochimica Acta, 167, 63–79.

Wieder, W. R., G. B. Bonan, and S. D. Allison (2013), Global soil carbon projections are improved by modelling microbial processes, Nature Climate Change, 3(10), 909–912.

Wieder, W. R., C. C. Cleveland, W. K. Smith, and K. Todd-Brown (2015), Future productivity and carbon storage limited by terrestrial nutrient availability, Nature Geoscience, 8(6), 441–444.

Willaman, J. J., and B. G. Schubert (1961), Alkaloid-bearing plants and their contained alkaloids, Agricultural Research Service, US Department of Agriculture.

Zachara, J. M., P. E. Long, J. Bargar, J. A. Davis, P. Fox, J. K. Fredrickson, M. D. Freshley, A. E. Konopka, C. Liu, and J. P. McKinley (2013), Persistence of uranium groundwater plumes: Contrasting mechanisms at two DOE sites in the groundwater–river interaction zone, Journal of contaminant hydrology, 147, 45–72.

Zhang, L., S. Wang, Y. Xu, Q. Shi, H. Zhao, B. Jiang, and J. Yang (2016), Molecular characterization of lake sediment WEON by Fourier transform ion cyclotron resonance mass spectrometry and its environmental implications, Water Research, 106, 196–203.

